# Access to PCNA by Srs2 and Elg1 controls the choice between alternative repair pathways in yeast

**DOI:** 10.1101/2020.03.24.006932

**Authors:** Matan Arbel, Alex Bronstein, Soumitra Sau, Batia Liefshitz, Martin Kupiec

**Affiliations:** School of Molecular Cell Biology and Biotechnology, Tel Aviv University, Ramat Aviv 69978

## Abstract

During DNA replication stalling can occur when the replicative DNA polymerases encounter lesions or hard-to replicate regions. Under these circumstances the processivity factor PCNA gets ubiquitylated at lysine 164, inducing the DNA damage tolerance (DDT) mechanisms that can bypass lesions encountered during DNA replication. PCNA can also be SUMOylated at the same residue or at lysine 127. Surprisingly, *pol30-K164R* mutants display a higher degree of sensitivity to DNA damaging agents than *pol30-KK127,164RR* strains, unable to modify any of the lysines. Here we show that in addition to trans-lesion synthesis and strand-transfer DTT mechanisms, an alternative repair mechanism (“salvage recombination”) that copies information from the sister chromatid, is repressed by the recruitment of Srs2 to SUMOylated PCNA. Overexpression of Elg1, the PCNA unloader, or of the recombination protein Rad52 allows its activation. We dissect the genetic requirements for this pathway, as well as the interactions between Srs2 and Elg1.

## INTRODUCTION

The integrity of the genome is very often compromised by internal and external sources of DNA damage. The vulnerability of the genome increases during DNA replication, when the DNA has to be unpacked and exposed (1, 2). Chemical modifications of the DNA or proteins bound to it can cause fork stalling or even collapse, leading to a situation in which DNA replication is not completed. To deal with this situation, complex cellular mechanisms have evolved. These response mechanisms act either to promote repair of the lesions or to allow their bypass, thus preventing them from being converted into fatal genomic rearrangements (3). The genetic pathways responsible for DNA repair and genome stability are highly conserved across species. One of these pathways, the DNA damage tolerance (DDT) pathway [also known as the *RAD6* or post-replication repair (PRR) pathway], is activated when single-stranded DNA (ssDNA) accumulates at stalled forks or at gaps created by re-priming downstream to the initial stalling lesion (4).

PCNA, the sliding clamp that acts as a processivity factor for replicative DNA polymerases, plays an important role in regulating the DDT. To this day, two main sub-pathways have been characterized. The first is activated by mono-ubiquitylation at lysine 164 of PCNA, and is mediated by the E2-conjugating enzyme Rad6 and the E3-ubiquitin ligase Rad18. This ubiquitylation takes place at sites of fork arrest where replication protein A (RPA) accumulates (5, 6) and promotes the exchange of the replicative DNA polymerase by a trans-lesion DNA polymerase able to bypass the DNA lesion by synthesizing DNA in an error-prone manner in most of the cases (7–10) (Trans Lesion Synthesis or TLS). Alternatively, the mono-ubiquitin can be extended by the E2 enzymes Ubc13–Mms2 (UBC13-UEV1 in mammals) together with the E3 Rad5 (or its mammalian orthologs, SHPRH and HLTF), to create K63-linked ubiquitin chains (11). This acts as a signal to direct damage bypass via a template switch involving the sister chromatid (Template Switch, TS)(12, 13). The exact nature of how the poly-ubiquitylation is recognized, and the molecular details of the bypass are still quite mysterious (14), although some form of sequence homology recognition is required (15, 16). In addition to ubiquitylation, PCNA can undergo SUMOylation, predominantly at lysine 164, and to a lesser extent, at lysine 127 (12). PCNA SUMOylation at K164 seems to be evolutionary conserved and has been observed also in other organisms (17, 18). SUMOylation of PCNA can lead to the recruitment of the helicase Srs2, a UvrD-like helicase that can disrupt Rad51 presynaptic filaments and thus prevent homologous recombination (HR) (19–23). Functional homologs of Srs2 seem to exist in other organisms [e.g.: PARI (18); RTEL1 (24)]. PCNA is loaded onto DNA by the RFC complex, and is unloaded by an alternative clamp unloader containing the Elg1 protein (25–27). The mammalian ortholog of Elg1, ATAD5, is also a PCNA unloader, important for genome stability and acts as a tumor-suppressing gene (28).

In addition to TLS and TS, damage can be resolved by a mechanism that involves HR proteins and is independent of PCNA ubiquitylation. This mechanism, hereafter referred to as ‘‘salvage recombination” (SR) is restrained by the Srs2 helicase (14, 29). This pathway is considered a last resort, as unchecked homologous recombination may generate genome instability (30). Although sometimes presented as three clear sub-pathways, the relationship between TLS, TS and SR are still mysterious, and proteins may be involved in more than one category: for example, Rad5, which is required to initiate the TS sub-pathway by poly-ubiquitylating PCNA, plays also a role in the recruitment of TLS polymerases (31). The dissection of the various branches is made even more complex by the variations in timing and location: damage bypass can take place at the fork, or in gaps left behind it by re-initiation; it can also occur in S-phase, during the actual replication, or later, in G2 (4, 32–34).

Here we analyze the role played by Srs2 in preventing the usage of the SR branch. We show that recruitment of Srs2 by SUMOylation of either K164 or K127 of PCNA abolishes its use. The SR sub-pathway can however be activated by overexpression of Elg1 or Rad52. The SR pathway requires Rad51, Rad52, Rad59, Sgs1 and Elg1 activities.

## RESULTS

### SRS2 may inhibit DDT by binding to K127 SUMOylated PCNA

To analyze the effect of mutating different genes, we performed quantitative serial dilution assays on a large number of MMS concentrations differing by small increments. This method allows determining the relative sensitivity of all the isogenic strains with high accuracy and consistency.

Mutation of lysine 164 on PCNA prevents this residue from undergoing ubiquitylation or SUMOylation. Lack of these modifications causes sensitivity to DNA damaging agents, due to the inactivation of the DDT pathways. PCNA can also undergo SUMOylation on lysine 127; however, mutants unable to carry out this modification are not sensitive to DNA damaging agents. Paradoxically the double mutant at lysines 127 and 164 *(pol30-KK127,164RR*, hereafter referred to as *pol30-RR)* is **<u>less</u>** sensitive to DNA damage than the single *K164R* mutant [(12)(Figure 1A)]. This finding is surprising and suggests that SUMOylation of PCNA at lysine 127 has an **<u>inhibitory</u>** effect on DNA damage repair or tolerance when K164 is mutated. SUMOylation of PCNA at K164 is coordinated by the Siz1 E3 ligase (35, 36); however, it is not clear which E3 ligase SUMOylates K127. When we deleted *SIZ1* in the background of *pol30-K164R* a partial suppression was observed. Since in yeast cells *SIZ2*, a paralog of *SIZ1*, can in some cases compensate for lack of Siz1 activity and to some extent promote K127 SUMOylation (37), we also combined *pol30-K164R* with *Δsiz2* or *Δsiz1 Δsiz2*. Although the single *Δsiz2* allele shows no effect, the *Δsiz1* or the *Δsiz1 Δsiz2* strains showed the same MMS resistance as *pol30-RR* cells (Figure 1A). Thus, Siz1 is the main enzyme involved, and inactivation of SUMOylation or mutation of K127 result in a similar effect. The reduced MMS sensitivity of *pol30-K164R* when there is no SUMOylation (*Δsiz1 Δsiz2 pol30-K164R* or *pol30-RR*) can be attributed to the lack of recruitment of Srs2 to PCNA by SUMOylation of lysine K127. Indeed, Figure 1B shows that mutating lysine 127 in a *pol30-K164R* strain *(pol30-RR)* has the same effect as deleting the *SRS2* gene, and that deleting *SRS2* in a *pol30-RR* strain has no further effect. We thus conclude that the recruitment of Srs2 to PCNA SUMOylated at K127 causes sensitivity to MMS when K164 is unmodified. Deletion of *SRS2* suppresses the high sensitivity to genotoxic agents of *Δrad18, Δrad5* and *pol30-K164* mutants, demonstrating that the binding of Srs2 to SUMOylated K127 has a major role in inhibiting repair (12).

**Figure 1:**
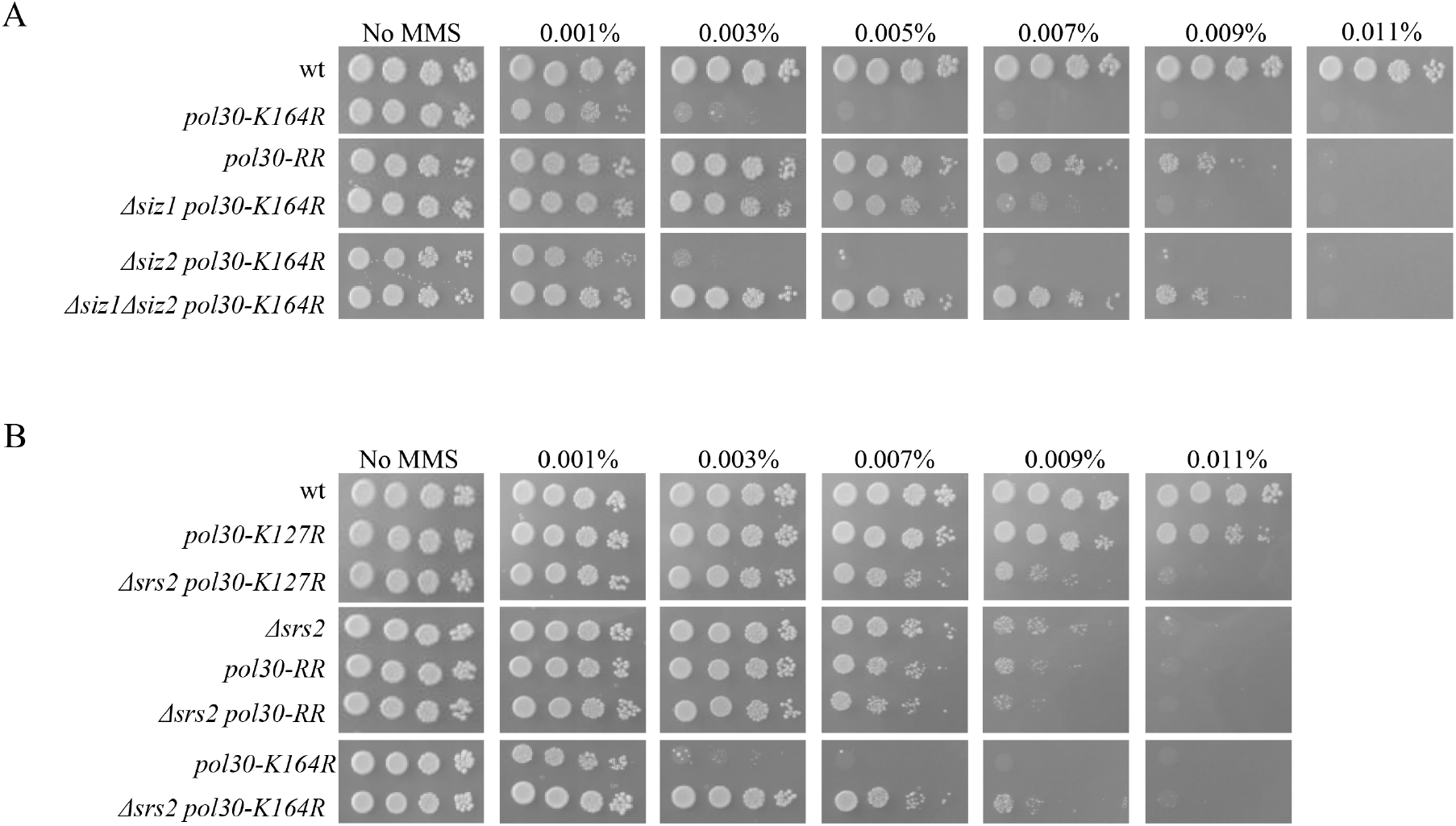
Srs2 and PCNA modifications show epistatic interactions. **(A)** Siz1 (and in its absence Siz2) has a role in SUMOylation of PCNA on lysine 127. **(B)** Deletion of *SRS2* exhibits complete epistasis to PCNA modification mutants. Ten-fold serial dilutions spotted on plates with increasing concentrations of MMS, photographed after 3 days.

### Overexpression of Rad52 and Elg1 suppress the sensitivity of *pol30-K164R* strains to MMS

The Srs2 helicase is able of evicting Rad51 from DNA and thus inhibiting DNA repair by homologous recombination (HR) (19, 20). It has thus been proposed that Srs2 bound to PCNA plays a role in preventing uncontrolled recombination events during the S phase. However, the exact mechanism by which Srs2 sensitizes mutants of the DDT pathway is still unclear.

In a *pol30-K164R* strain Srs2 can bind only to K127; yet, the helicase still plays a negative role, as evidenced by the fact that deletion of *SRS2* restores resistance to MMS (Figure 1B). In order to better understand the mechanism by which Srs2 exerts its negative effect, we asked whether it was possible to bypass its effect by overexpressing members of the HR machinery.

Figure 2A shows that overexpression (OE) of Rad52 can partly suppress the sensitivity of *pol30-K164R* to MMS. A weak suppression is also seen by OE of Rad54, but not by Rad51, Rad55, Rad57 or Rad59 (Figure 2A). The fact that Rad51 had no suppression effect was very surprising, as this recombinase is central to most HR-related biochemical reactions, and Srs2 is known to inhibit its activity. In addition, co-overexpression of Rad51+Rad52 gave no further suppression than that provided by Rad52 (Figure 2A). Therefore, Rad51 protein levels are not limiting for the suppression effect.

**Figure 2:**
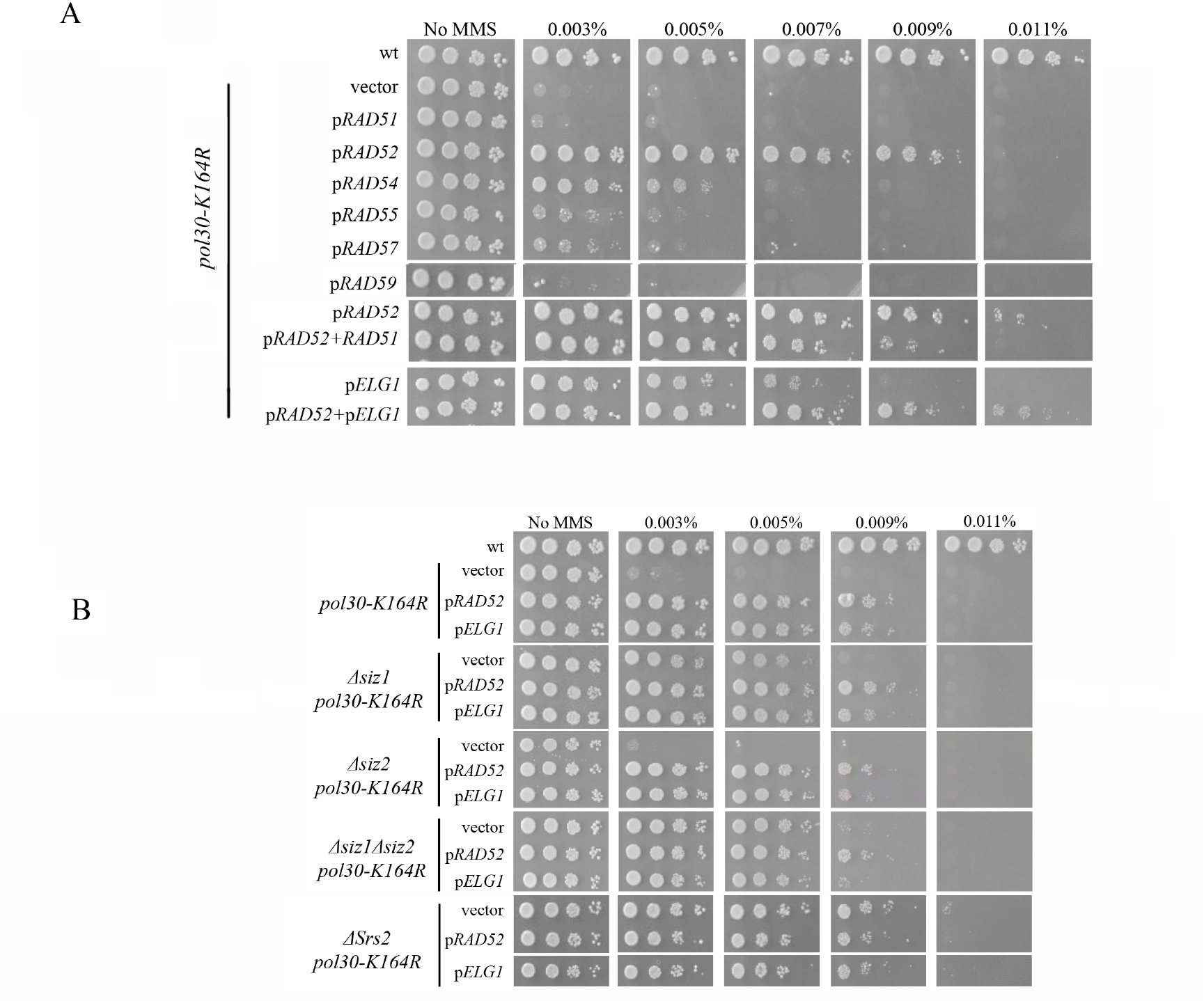
Suppression of MMS sensitivity by overexpression (OE) of Rad52 or Elg1. *pol30-K164R* strains were transformed with various high-copy number plasmids. **(A)** OE of Rad52 and Elg1, but not of Rad51 or Rad59, can suppress the sensitivity o f*pol30-K164R* to MMS. Elg1 and Rad52 overexpression act in the same pathway. **(B)** Overexpression of Rad52 or Elg1 are epistatic to *Δsiz1, Δsiz2, Δsiz1Δsiz2* and *Δsrs2* in the *pol30-K164R* background.

In addition to Srs2, Elg1, the unloader of PCNA, also interacts with SUMOylated PCNA (25). By competing for the same binding sites on PCNA, or unloading PCNA altogether, Elg1 may potentially limit the access of Srs2 to the DNA and lower its activity. Overexpression of Elg1 suppresses the MMS sensitivity of *pol30-K164R* almost as well as overexpression of Rad52 (Figure 2A). We next asked whether *RAD52* and *ELG1* work together or in separate pathways. Overexpressing the two genes together had no additive effect (Figure 2A), suggesting that they work by a common mechanism.

### Overexpression of Rad52 or of Elg1 allow increased damage tolerance by bypassing Srs2 inhibition

Mutant *pol30-K164R* strains overexpressing Rad52 or Elg1 show a suppression level similar to the one observed in *pol30-K164R* strains carrying the *Δsiz1Δsiz2* or the *Δsrs2* mutations. To test whether different pathways are involved, we overexpressed either Rad52 or Elg1 in *pol30-K164R Δsiz1Δsiz2* (no SUMOylation of PCNA at lysine 127) and *pol30-K164R Δsrs2* (no Srs2 recruitment) strains. No increased MMS resistance was observed, suggesting that OE of Rad52/Elg1 suppresses *pol30-K164R* sensitivity by acting in the same pathway as that affected by lack of K127 SUMOylation or deletion of Srs2 (Figure 2B). Interestingly, overexpression of Rad52 or Elg1 resulted in the same level of sensitivity to MMS in *pol30-K164R Δsiz1, pol30-K164R Δsiz2* and *pol30-K164R Δsiz1 Δsiz2* strains, despite their different sensitivity levels in the absence of the overexpressing plasmids (Figure 2B). Taken together, these results hint at a common suppression mechanism for Srs2 depletion from the site of DNA damage and for Rad52 or Elg1 overexpression.

To better understand the genetic connection between Rad52 and Elg1, a genetic dependency analysis was carried out. Figure 3A shows that overexpression of either *RAD52* or *ELG1* fails to suppress the *pol30-K164R* sensitivity in the absence of the *RAD51* gene. This suggests that (despite the fact that Rad51 OE had no effect), a Rad51-mediated mechanism is needed for the suppression observed. Moreover, suppression of *pol30-K164R* by *ELG1* OE requires *RAD52*, and suppression by *RAD52* OE is partially dependent of *ELG1*. These results are in line with our previous observation, showing that both genes work in the same pathway (Figure 2A). Deletion of *RAD59* partly abolished the suppression of *pol30-K164R* by either *RAD52* or *ELG1* OE. Rad59 is a Rad52 paralog that lacks Rad51-interacting regions and becomes important for DNA repair in the absence of Rad51 (38, 39). Our results show that Rad59 is needed for the efficient work of Rad52 in this pathway, although its protein levels are not limiting, and thus its overexpression has no effect (Figure 3A). The Salvage Recombination pathway is expected to create sister chromatid junctions (SCJs), recombination structures that display exchange of strands between sister chromatids. These structures are eventually resolved by the activity of the Sgs1 helicase (40–42). Indeed, deletion of the *SGS1* gene abolished the suppression of *pol30-K164R* by either *RAD52* or *ELG1* OE (Figure 3B).

**Figure 3:**
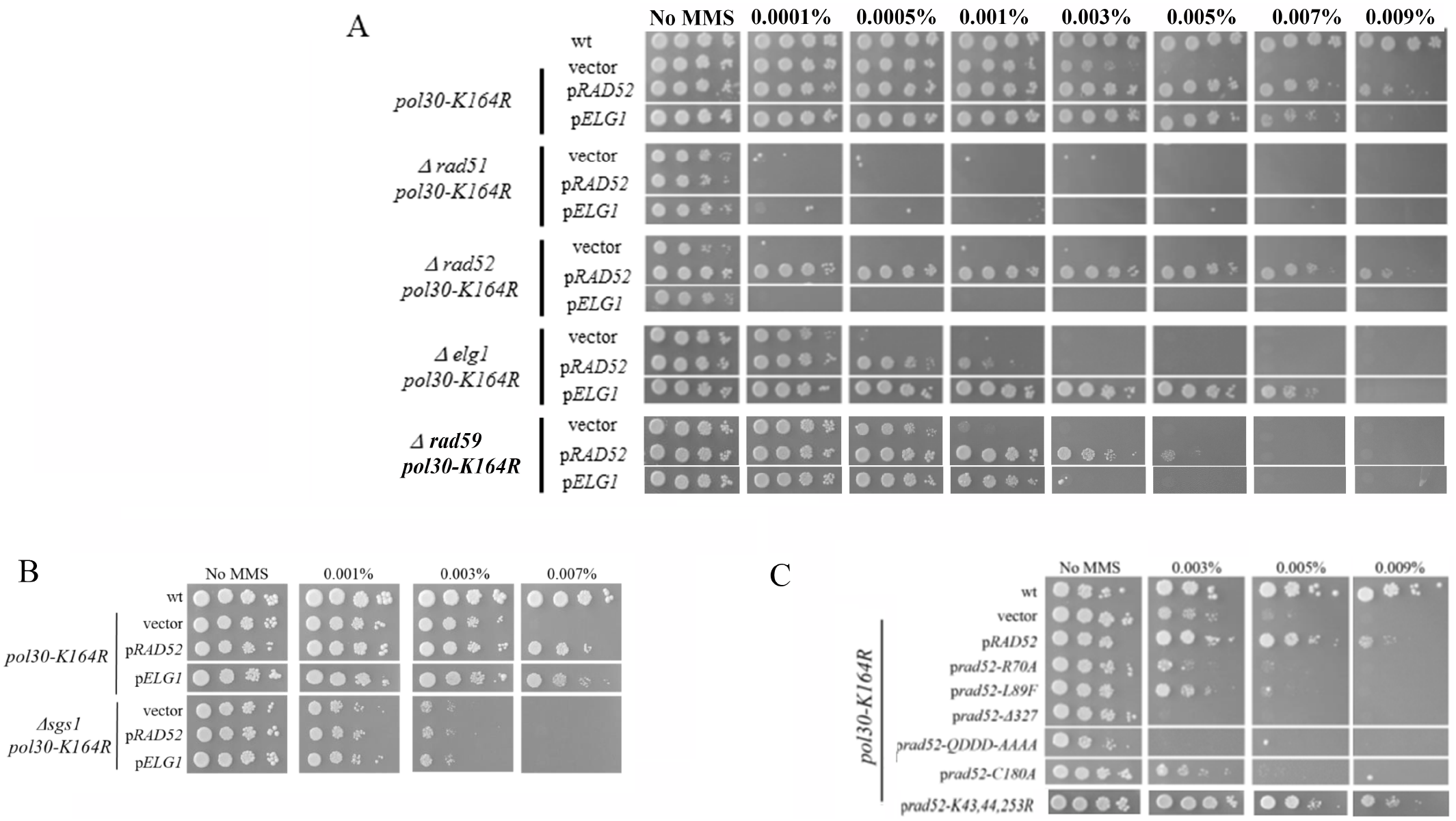
**(A)** The effect of overexpressing Rad52 and Elg1 on *pol30-K164R* strains require Rad51, Rad52, Rad59 and Elg1. **(B)** The effect of overexpressing Rad52 and Elg1 on *pol30-K164R* strains require Sgs1. **(C)** Analysis of Rad52’s motifs required for the suppression effect. A *pol30-K164R* strain was transformed with high copy number plasmids carrying *RAD52* alleles. See text for description of each allele.

In summary, our results suggest a model in which the repair pathway requires the removal of PCNA, a process carried out mainly by Elg1. This in turn, allows the Rad52-Rad59 complex to initiate the repair on the damaged site, by annealing to the sister chromatid. Finally, the SCJs thus created are resolved by Sgs1. The partial requirement for Elg1in the SR pathway (compared to the essentiality of Rad52) can be accounted by the fact that alternative mechanisms of PCNA unloading exist (and thus, despite its important role during DNA replication, *ELG1* is not an essential gene).

To further analyse the role of Rad52 in the suppression effect, we overexpressed, in the *pol30-K164R* strain, Rad52 mutants defective in specific functions of Rad52, and tested their suppression ability. The *rad52-R70A* and *rad52-L89F* mutants are unable to perform the DNA annealing function of Rad52 (43),(44). Both mutants are unable to suppress *pol30-K164R* (Figure 3C), implying that the suppression effect acts through the Rad52 DNA annealing ability. The *rad52-Δ327* allele (45) lacks the C-ter of Rad52, which is required for interaction with Rad51. The *rad52-QDDD-AAAA* allele (45) is impaired in its interaction with RPA and recombination mediator activity. Both alleles are thus unable to stimulate Rad51 activities. When overexpressed in a *pol30-K164R* strain, they failed to suppress, and instead sensitized cells to MMS (Figure 3C). This again is consistent with a role for Rad51 in the Rad52 OE-stimulated repair of lesions. The *rad52-C180A* allele was described as unable to utilize sister chromatid information for repair (46), and, as expected, also exhibited no suppression effect (Figure 3C). Rad52 undergoes SUMOylation after DNA damage, and lack of SUMOylation affects its stability but does not affect its recruitment to DNA damage (27, 28). However, Rad52 SUMOylation may also disturb Rad51 filament formation by the activity of the SUMO-targeted Cdc48 segregase, which can limit Rad52-Rad51 physical interaction and displace them from DNA (29, 30). When a *RAD52* allele unable to undergo SUMOylation *(rad52-K43,44, 253R)* was overexpressed, it was still as potent a suppressor of the *pol30-K164R* allele (Figure 3C). We thus conclude that lack of SUMOylation on Rad52 does not affect its suppression capabilities.

The analysis of *rad52* mutants thus suggests that Rad52-mediated sister-chromatid-dependent repair requires the DNA annealing ability of Rad52, as well as its function in promoting assembly of Rad51 nucleofilaments, but not its SUMOylation.

### Rad52 and Rad51 interact with the same region of Srs2

Our results up to now suggest that in *pol30-K164R* strains, Rad52 and Elg1 OE work in the same SR sub-pathway, which includes DNA annealing with the sister chromatid, and bypasses the restraint exerted by Srs2 recruited to SUMOylated K127 of PCNA. OE of Rad52 may suppress the sensitivity to DNA damaging agents by facilitating the access of Rad52 to the DNA damage site even when Srs2 is present; Elg1 activity unloads PCNA, and thus may reduce the level of Srs2 recruitment and its negative effects. Of note, the inhibition of repair by Srs2 is not likely to be by Rad51 eviction, since OE of Rad51, in contrast to that of Rad52, has no suppression effect. Instead, we assume that Srs2 directly inhibits, or counteracts, the activity of Rad52 (21, 22). In support of this idea, a physical interaction was characterized between Srs2 and Rad52 (47).

We preformed yeast two hybrid experiments in which plasmid containing Rad52 or Rad51 fused with the transcription activating domain of Gal4 and a plasmid containing Srs2 fused with the DNA binding domain of Gal4 were both transformed to a histidine auxotroph strain in which the *HIS3* gene has a Gal4-activated promoter. Growth on plates lacking histidine indicates that the two protein have physical interaction. Our own yeast two hybrid results identified the interaction between Srs2 and Rad52 in a *Δrad51* strain, again implying that this interaction is independent on Rad51 (Figure 4A). Furthermore, we found that Srs2 binds both Rad51 and Rad52. To pinpoint the exact interaction region of Srs2 and Rad52, we created a set of Y2H constructs carrying various regions of Srs2. Figure 4B shows that a fragment containing amino acids 875 to 902 (the same region that binds Rad51) binds Rad52. In contrast, either the N-terminal region alone, the C-terminal region alone, or the full fragment lacking the Rad51-interacting sequence (875–902), fail to show interaction in the Y2H assay (Figure 4A). Thus, Srs2-Rad51 and Srs2-Rad52 interactions are mediated by the same region of Srs2, and are independent of PCNA- and SUMO-interacting motifs (PIM and SIM, respectively).

**Figure 4:**
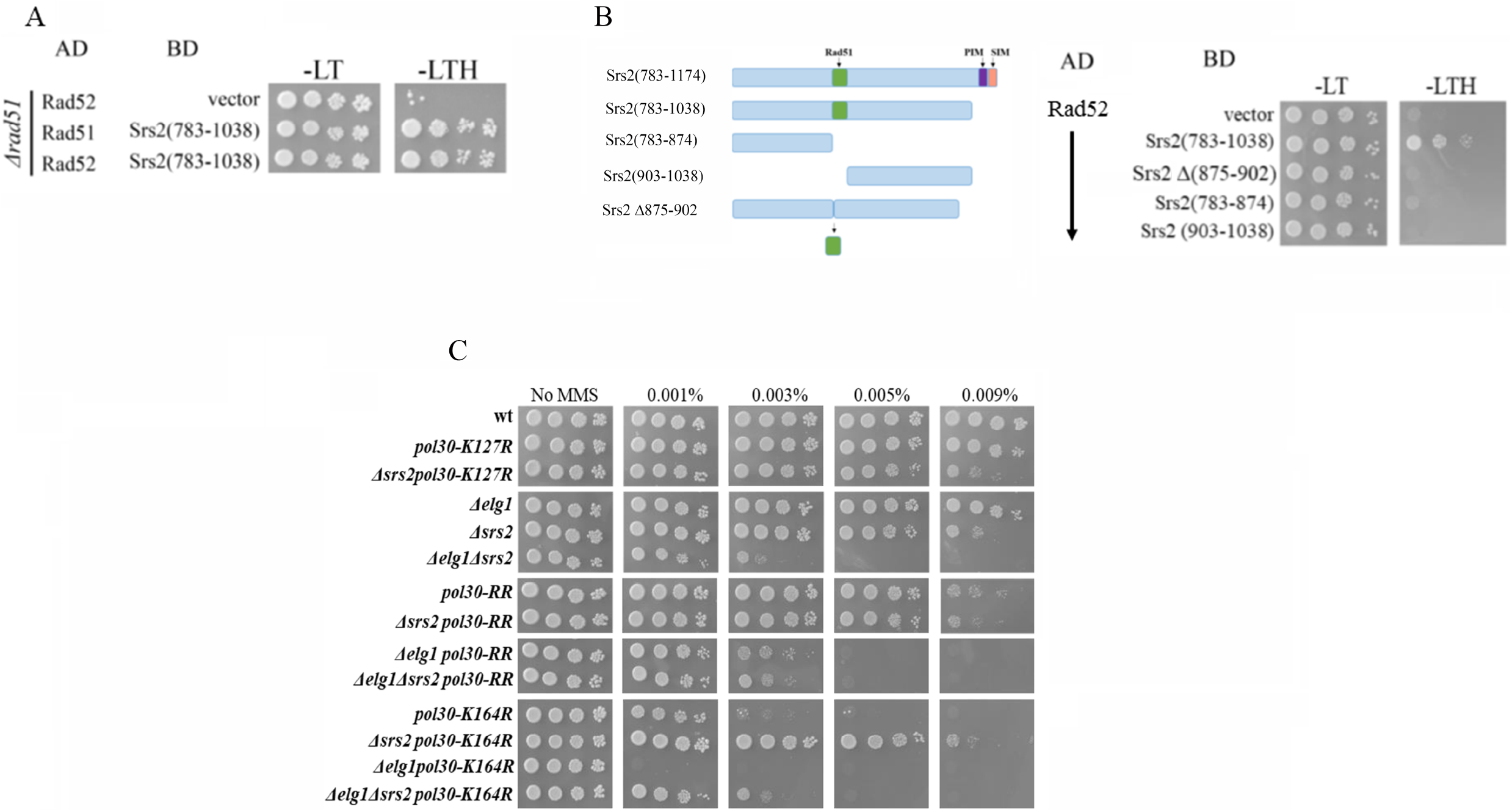
**(A)** Yeast-2-Hybrid experiment showing that Rad51 and Rad52 interact with the same region of Srs2, even in the absence of genomic *RAD51*. Plasmids containing either Rad52 or Rad51 fused with the transcription activating domain of Gal4 and a plasmid containing Srs2 fused with the DNA binding domain of Gal4 were transformed to the yeast strain with leucine and tryptophan selection respectively. The *HIS3* gene is under a Gal4-dependent promoter and thus was expressed only if physical interactions occur between the two expressed fusion proteins. –LT: medium lacking leucine and tryptophan; -LTH: medium lacking leucine, tryptophan and histidine. **(B)** Yeast-2-Hybrid experiment showing that Rad52 interacts exclusively with the region of Srs2 between amino acids 875 and 902 and independently of PCNA interacting motif and SUMO interacting motifs (PIM and SIM respectively). **(C)** Genetic interactions between *Δelg1, Δsrs2* and PCNA mutants.

### The interplay between Srs2 and Elg1 in the SR pathway

What is the relationship between Srs2 and Elg1? The simplest model would imply the following scenario: upon fork stalling, Srs2 is recruited to SUMOylated PCNA, repressing SR. Later, the Elg1 RLC unloads it together with the clamp. This model predicts that *Δsrs2* and *Δelg1* should show an epistatic relationship. However, this is not the case: the double mutant *Δsrs2 Δelg1* is clearly more sensitive than the single mutants [(48) and Figure 4C]. How can we explain this phenotype? As expected from our working model, deleting *SRS2* in a *pol30-K164R* strain results in reduced sensitivity to DNA damage, due to the opening of the salvage recombination pathway. Consistent with our results (Figure 3A), this sub-pathway relies on PCNA removal from the chromatin, which is done mainly by Elg1. Deleting *ELG1* in the double mutant *Δsrs2 pol30-K164R* reduces the sensitivity to the same level seen in *Δelg1 pol30-RR* (and *Δelg1 Δsrs2 pol30-RR):* in these strains, the TS and TLS branches are blocked by mutation of lysine 164 and the SR path is partially blocked by lack of Elg1 (Figure 4C).

Interestingly, the double mutant *Δelg1 pol30-K164R* (in which Srs2 is still active and **<u>can</u>** be recruited to K127, but the SR sub-pathway is blocked by the *Δelg1* mutation) shows a level of sensitivity higher than any of the double- and triple-mutants in which Srs2 recruitment to K127 of PCNA is blocked *(Δelg1 Δsrs2 pol30-K164R., Δelg1 pol30-RR Δelg1 Δsrs2 pol30-RR)*. These results suggest that even in the absence of Elg1, if Srs2 is not recruited, some repair is carried out. This is consistent with the fact that alternative PCNA unloading mechanisms exist, and that deletion of *ELG1* only partially blocks the SR pathway when Rad52 is overexpressed (Figure 3A).

### Srs2 recruited to SUMOylated K164 has a stronger effect on the SR pathway

Our results are consistent with a model in which SUMOylation on K127 allows the binding of Srs2, which restricts the activity Rad52. Next, we compared the recruitment of Srs2 at K127 with its recruitment at K164, the main amino acid of PCNA that undergoes SUMOylation.

First, we validated that *Δrad18* and *Δrad5*, responsible of mono- and poly-ubiquitylation at K164 of PCNA, are not required for the suppression effect of OE Rad52/Elg1. As expected, when Rad52 or Elg1 were overexpressed in *Δrad18 pol30-K164R* or *Δrad5 pol30-K164R* strains, they still showed suppression (Figure 5A).

In *Δrad18* or *Δrad5* strains harboring a wt PCNA, the TS and TLS branches in the DDT pathway are inactive, but K164 (in addition to K127) can still be SUMOylated, and thus is still able to recruit Srs2. Figure 5B shows that OE of *RAD52* in the background of *Δrad18* or *Δrad5* still causes suppression, albeit weaker than that seen in the *pol30-K164R* background. Interestingly, in these strains, joint OE of Rad52 and Rad51 shows an **<u>additive</u>** suppression effect that was not seen in *pol30-K164R*. These results imply that Srs2 bound to SUMOylated PCNA at K164 has a stronger negative effect on the SR sub-pathway, probably by evicting Rad51 from the DNA (19, 20). Overexpression of Rad51, in addition to Rad52, is needed to overcome this negative effect.

In the absence of functional *RAD18* or *RAD5*, when the DTT pathway is inactive, but Srs2 is recruited (mainly to K164), OE of Elg1 completely failed to show any suppressing effect (Figure 5B) and it had no additive effect with OE of Rad51 (data not shown). We conclude that Srs2 recruited by SUMOylated PCNA at K164 prevents Elg1 from unloading PCNA.

If recruited Srs2 exerts an inhibitory effect on Rad52 and Elg1, then we expect that deletion of *SRS2* in *Δrad18/Δrad5* strains will suppress their sensitivity (49, 50) (making them as resistant to DNA damage as *Δsrs2* alone [as shown for *pol30K164R* (Figure 1B)]. In figure 5C we see that this is the case, and that the cells are able to grow in MMS concentrations as high as 0.09% MMS, as do *Δsrs2* cells. As expected, the suppression depends on Elg1 activity, and *Δrad18/Δrad5* strains deleted for both *SRS2* and *ELG1* become sensitized again to MMS.

## DISCUSSION

### Recruitment of Srs2 by PCNA SUMOylation at either K127 or K164 controls the salvage recombination (SR) pathway

The error-free DDT plays a central role in dealing with stalled forks during DNA replication. However, evidence for a Rad18 and Rad5-independent recombinational repair mechanism, which uses information from the sister chromatid to allow repair of damage sites, has been documented and was termed the salvage pathway (15, 16, 51, 52). This mechanism requires Rad51 and Rad52 to invade the sister chromatid DNA, and Sgs1 to resolve the joint molecules crated by the template switch (14).

The role for Srs2 in regulating the salvage pathway has been deduced from studies showing that deletion of *SRS2*, similarly to *Δsiz1*, can suppress the DNA damage sensitivity of DDT mutants. The suppression depends on the interaction of Srs2 with SUMOylated PCNA (49, 50). However, the precise interplay between the factors that participate in the DDT and the salvage pathway are not well defined. Moreover, as most SUMOylation events are at K164, the role of K127 SUMOylation has not been described. In this work, we define a role for K127 SUMOylation and characterize the way in which it regulates the SR pathway. By analyzing the suppression level of Srs2, Siz1 and Siz2 in a background of Pol30 mutated in its lysine 164, we found that SUMOylation of Lysine 127 is carried out mainly by Siz1. Deletion of *SRS2* suppresses the MMS sensitivity of all the DDT mutants to the same level as *pol30-RR* (Figure 1B). Although we present results obtained with MMS, similar results were seen with other DNA damaging agents (data not shown). This suggests that Srs2 has full control over the SR pathway and must be removed from SUMOylated PCNA as a perquisite for the activity of the pathway. Following the removal of Srs2, PCNA must be removed [either by Elg1 or by alternative mechanism (25, 27, 53)], to allow repair.

### Elg1 and Rad52 promote the initiation of the salvage pathway

OE of Elg1 and Rad52 can suppress the MMS sensitivity of *pol30-K164R* to the same extent as deletion of *SRS2* (Figure 2B). Further examination revealed that OE of Rad51 has no suppression phenotype by itself, despite the fact that the suppression is **<u>dependent</u>** on Rad51 (compare Fig. 2A (pRAD51) with Fig. 3A (Δrad51 pol30k164R). Interestingly, on the background of *Δrad18* or *Δrad5* Rad51 OE has a phenotype when overexpressed together with Rad52 (Figure 5B). This observation, taken together with the fact that Y2H experiments show a direct interaction between Srs2 and Rad52, even in the absence of Rad51 (Figure 4A,B) shows that Srs2 represses the salvage pathway by directly inhibiting not only Rad51, but also Rad52. Similarly, in the background of *pol30-K164R*, the suppression effect of deleting *SRS2* depends on Elg1 (Figure 5C), stressing again the important role of Elg1 in SR. We have previously shown that in the absence of Elg1 there is an accumulation of SUMOylated PCNA, and Srs2, on the chromatin (25). Taking all these results in consideration, we propose that Srs2 and Elg1 compete for the interaction with PCNA. Overexpression of Elg1 thus removes the inhibition of Srs2 on SR, either because of direct competition, or because higher PCNA unloading activity by Elg1 leads to lower recruitment of Srs2. As Rad52 is a direct target, it is also possible to overcome the inhibition by overexpressing Rad52. Our results thus clarify the interplay between the various options for repair and lesion bypass. A model explaining the different phenotypes is further elaborated below (Figure 6).

**Figure 5:**
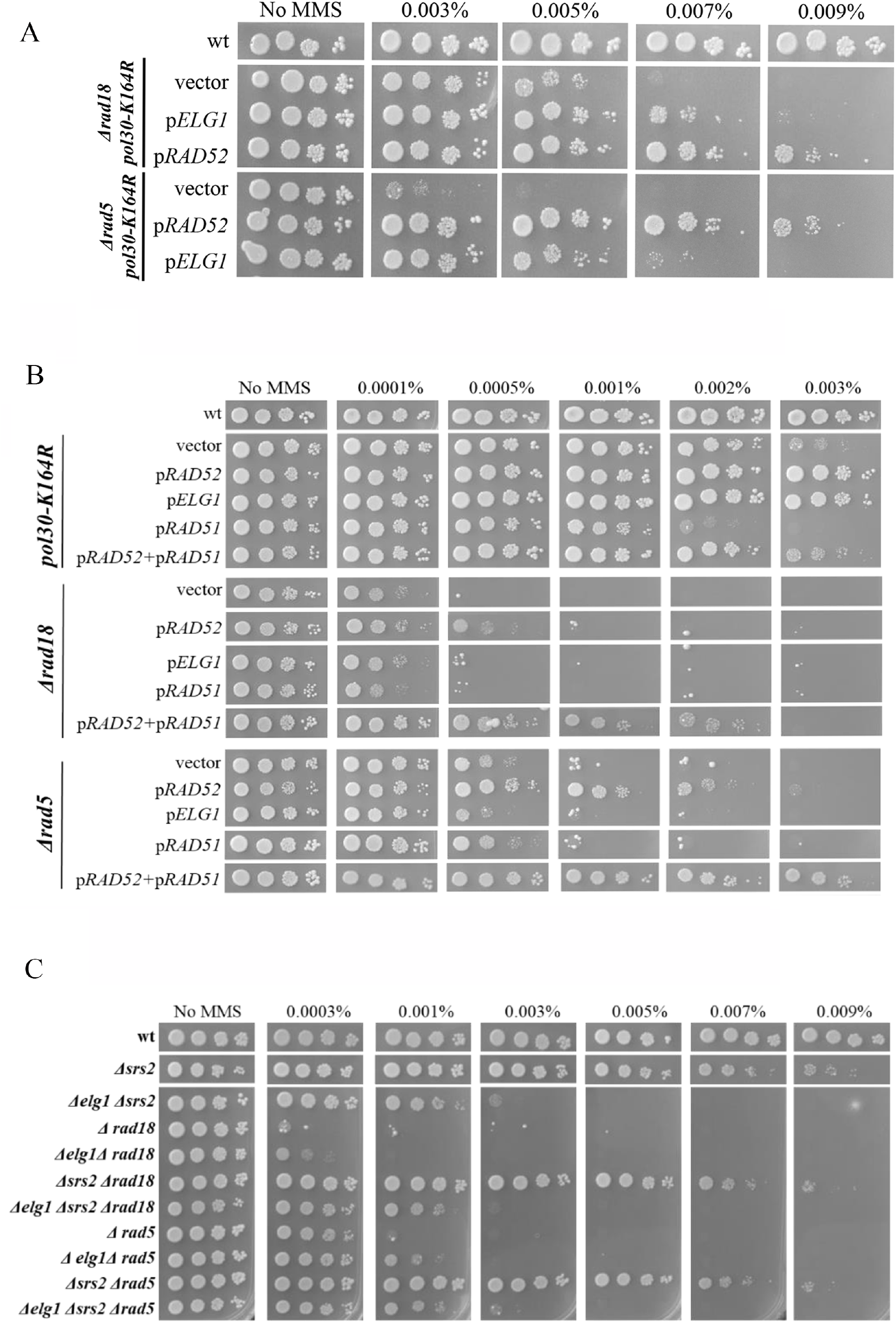
SUMOylation of K164 on PCNA has a role in the regulation of the repair pathways. **(A)** The suppression effect of *pol30-K164R* by OE of Rad52/Elg1 is still evident when Rad18 or Rad5 are not available. **(B)** OE of Rad52/Elg1 is less effective when the DDT pathways are inactive but K164 can bind Srs2. **(C)** Srs2 deletion suppresses the DDT defective mutants to the same level.

**Figure 6:**
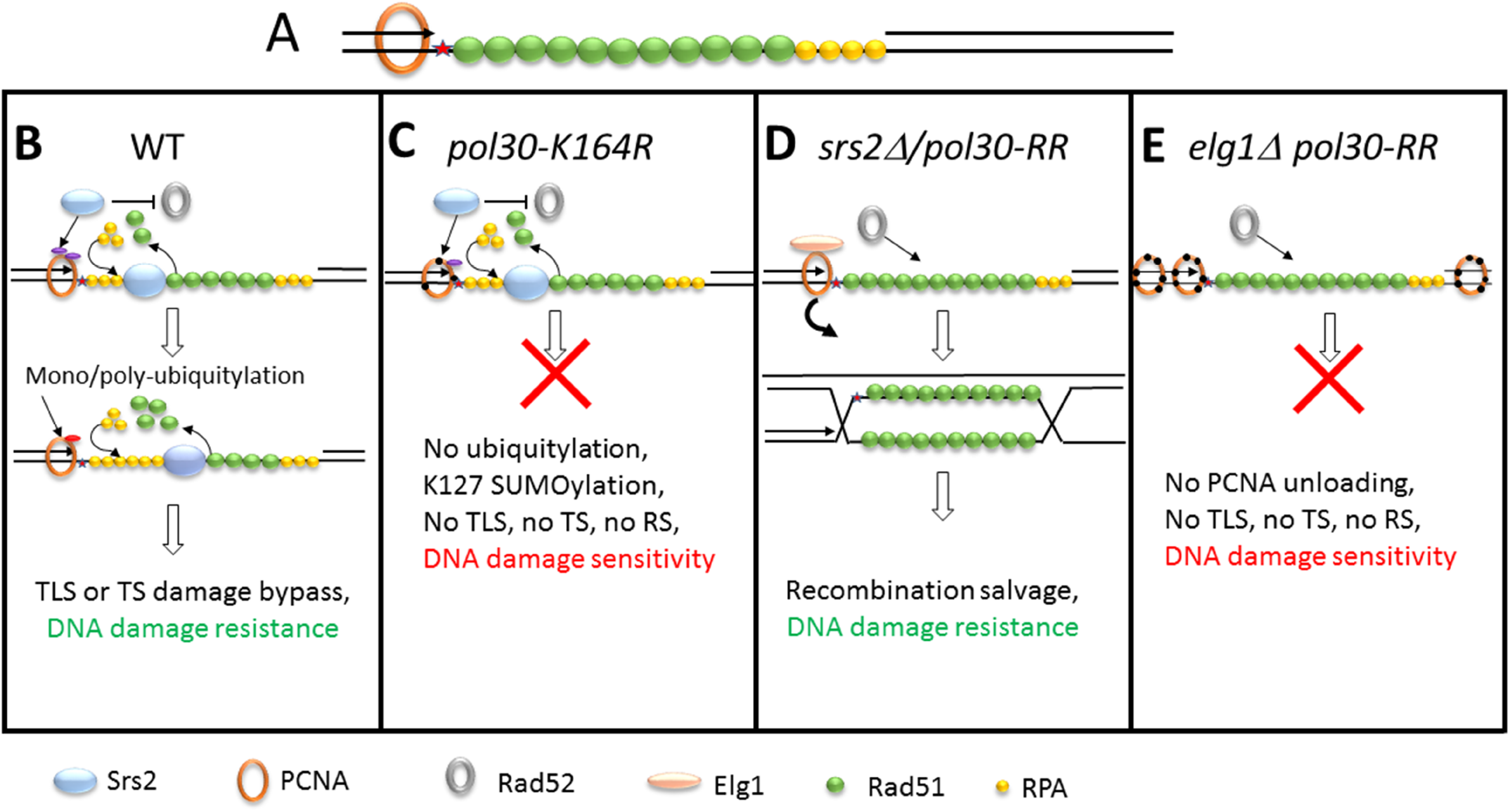
Schematic representation of the recombination salvage pathway regulation. **(A)** Lesions or perturbations in the DNA (red star) cause fork arrest. The model presented here can take place at the stalled replication fork, or, as shown, at ssDNA gaps left behind by arrested replication followed by re-start downstream. Events can occur during DNA replication (S-phase) or after the bulk of the replication has been completed (G2 phase). ssDNA will be immediately covered by RPA and Rad51. **(B)** During normal DNA replication Srs2 binds to both SUMOylated lysines of PCNA. It inhibits the activity of Rad52 and evict Rad51, thus precluding homologous recombination events. PCNA ubiquitylation will lead to damage bypass by translesion synthesis (TLS) or by template switch (TS). The choice between these two sub-pathways may depend on the nature and amount of accumulated DNA damage. **(C)** When lysine 164 of PCNA is mutated, Srs2 is still recruited by K127 SUMOylation. Srs2 inhibits the activity of rad52 and Rad51, and since PCNA cannot be ubiquitylated, no DDT sub-pathway is available. The cells are thus extremely sensitive to DNA damage. **(D)** When Srs2 is deleted, or both lysines of PCNA that participate in Srs2 recruitment are mutated *(pol30-RR)* the salvage recombination is driven by Rad52 and Rad51. This requires unloading of PCNA by the Elg1 RFC-like complex (although some level of leakage exists). **(E)** In a *Δelg1 pol30-RR* strain no TS or TLS recruitment is possible (as PCNA cannot get modified), and Srs2 is not recruited; however, the SR pathway cannot proceed because PCNA cannot be efficiently unloaded. As a consequence, the cells are sensitive to DNA damage.

### SUMOylated lysine 164 is the main recruitment site for Srs2

Analysis of the interplay between the two SUMO sites of PCNA when the DDT pathway is not active, reveals that deletion of *SRS2* or mutation of lysines 127 and 164 in PCNA suppresses the MMS sensitivity of *Δrad5* or *Δrad18* mutants. This suppression depends on the activity of Elg1 (Figures 3A and 5C). Unlike Rad52 OE, Elg1 OE did not show any suppression in *Δrad5* or *Δrad18* cells (Figure 5B), which are still able to undergo full SUMOylation of PCNA. Furthermore, OE of Rad51 with Rad52 showed **<u>additive</u>** effects in the background of *Δrad5* or *Δrad18*, but not in *pol30-K164R* (Figures 5B and 2A). We interpret these results as follows: SUMOylated lysine 164 is the major recruiter of Srs2. Srs2 bound to SUMOylated lysine 164 regulates the salvage pathway by inhibiting the activities of Rad52 and Rad51. In contrast, recruitment of Srs2 to the SUMOylated lysine 127 of Pol30 is rarer (SUMOylation on lysine 127 can be detected at a much lower levels than on lysine 164). Lower recruitment of Srs2 on the DNA damage site may allow sporadic SR events to take place, explaining the positive OE phenotype of Elg1 in K164R but not in *Δrad5* or *Δrad18*. Thus, Srs2 inhibition on lysine 164 of PCNA has a more general role in controlling SR, as opposed to recruitment through lysine 127, which may be time- and place-specific. Recent experiments have shown, for example, that repair and checkpoint signaling differ at ssDNA gaps and at stalled forks (33). PCNA modifications may be different at these two locations. An alternative, not necessarily exclusive possibility is that Srs2 bound to K127 acts by inhibiting Rad52, whereas binding to the K164 residue of PCNA activates Srs2’s Rad51 evicting activity. Our results could perfectly fit such a scenario.

### A model for the salvage pathway regulation

Taking all the obtained results together, we propose the following model (Figure 6): During normal S-phase progression, PCNA at arrested forks, or left behind at ssDNA gaps (Figure 6A) undergoes SUMOylation, and Srs2 can bind to either SUMOylated lysine of PCNA (mainly K164). Srs2 inhibits the activity of Rad52, Rad51 and Elg1, thus preventing untimely/unwanted recombination events. Increased levels of RPA near the arrested PCNA molecule allow it to become mono-ubiquitylated by the Rad6/Rad18 complex (6), and, if necessary, further poly-ubiquitylated by the Rad5/Mms2/Ubc13 complex to allow TS (Figure 6B). The mechanism that decides between these two branches remains mysterious. In *pol30-K164R* strains, Srs2 is still recruited by SUMO at K127, and thus the SR option is closed, but ubiquitylation of PCNA is precluded by the mutation at K164. This results in increased sensitivity to DNA damage (Figure 1A and Figure 6C). In the absence of Srs2, or when PCNA cannot be modified *(pol30-RR)* cells are **<u>less</u>** sensitive because the SR pathway is open: this requires unloading of PCNA (mainly by Elg1) to invade the sister chromatid, as well as the Rad51, Rad52, Rad59 and Sgs1 proteins (Figure 6D). Finally, deleting *ELG1* in the *pol30-RR* strain leaves the cells with only partially active SR option, and thus more sensitive to DNA damage (Figure 6E).

Our results shed light on the important role that PCNA and its modifications have in determining the type of lesion bypass/repair used by the cell, and show how complex these decisions are. They also better define the role of Srs2 in the repair of damage during DNA replication. In this study, we have used a minimalist approach, and left out additional levels of regulation, such as Srs2 modifications and cellular regulators. Additional players that have an impact on pathway choice are Uls1 (54), Esc2 (29), Mph1/Mhf1-2 (55, 56), and others. Future work will center on the role played by these proteins, as well as the choice between TS and TLS, both of which require PCNA ubiquitylation.

## Materials and Methods

### Yeast strains

Unless differently stated, all strains are derivatives of MK166:

### MK166

*MATa lys2:: Ty1Sup ade2-1(o) can1-100(o) ura3-52 leu2-3, 112 his3del200 trp1del901 HIS3:: lys2:: ura3 his4:: TRP1:: his4*. (57)

Standard Yeast Molecular genetics techniques were used to delete individual genes.

Strains and plasmids list are available in Table 1 and Table 2 respectively.

**Table 1:**
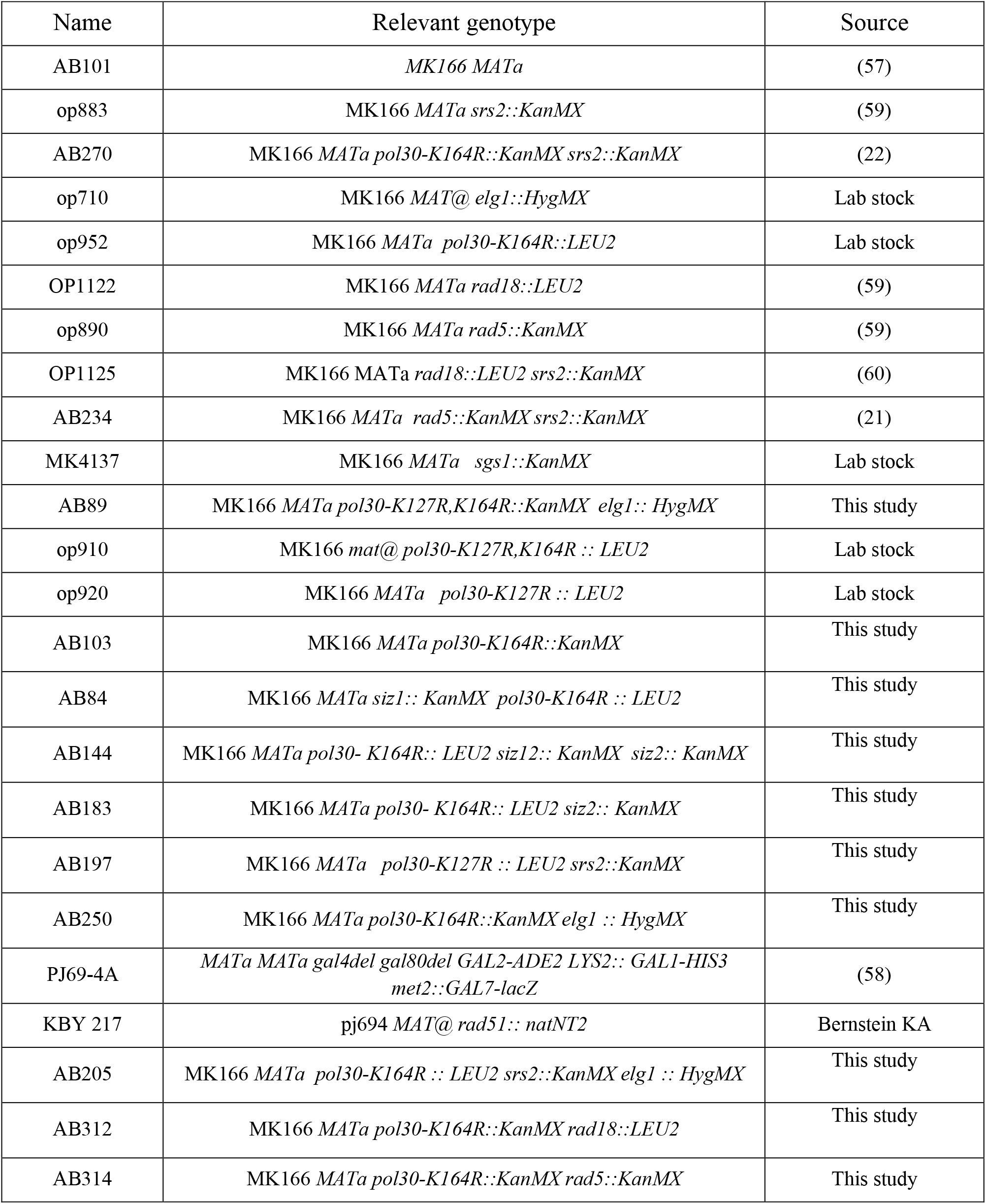
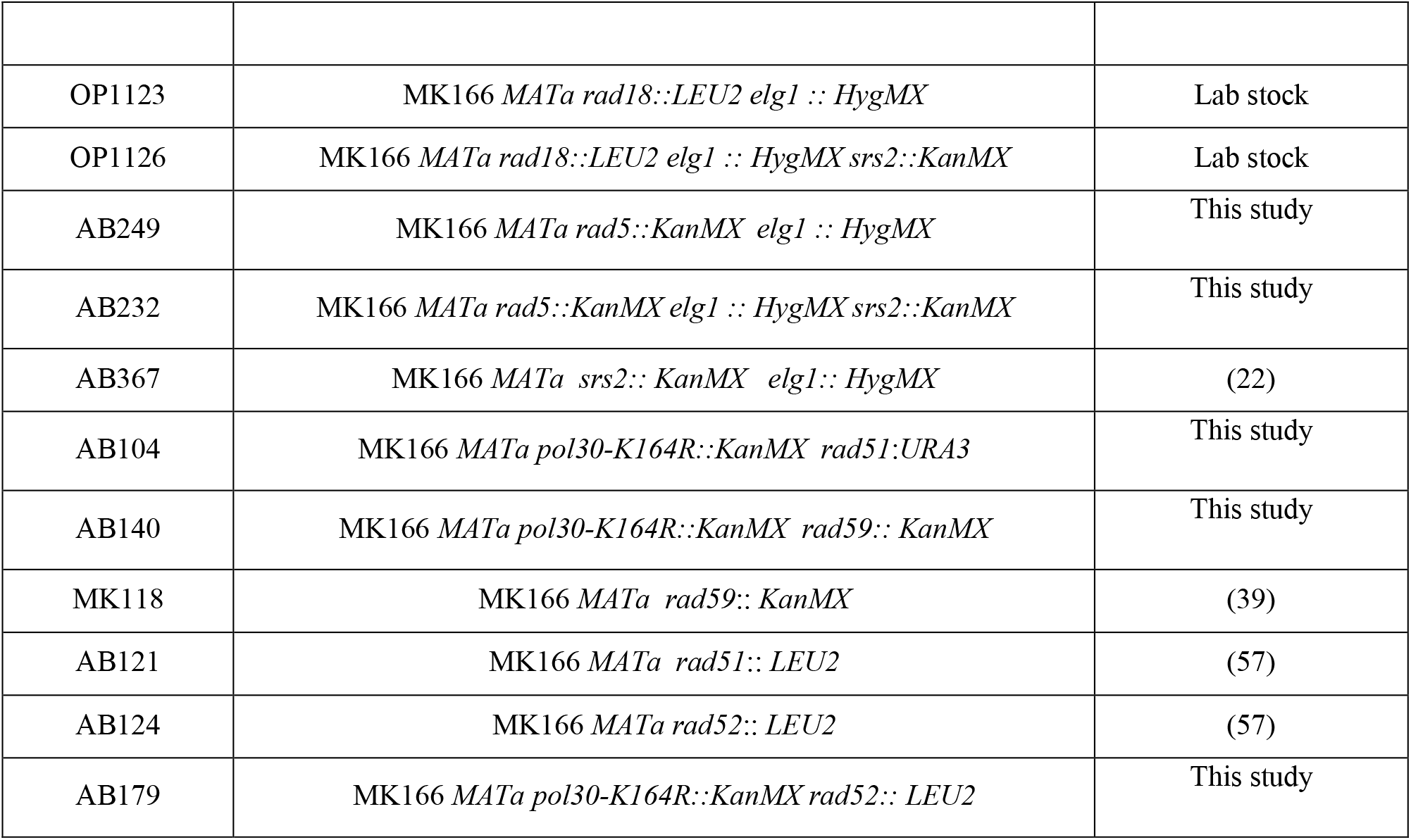
Yeast strain list

**Table 2:** Plasmid list

| Name | Relevant genotype | Source |
| --- | --- | --- |
| WDH262 | *RAD57* in 2 micron *LEU2* | David Schild |
| pYEp13-RAD54 | *RAD54* in 2 micron *LEU2* | David Schild |
| WDH261 | *RAD52* in 2 micron *LEU2* | David Schild |
| AB067B | *RAD52* in pRS426 | Lorraine Symington |
| WDH264 | *RAD55* in YEp13 | Wolf-Dietrich Heyer |
| WDH265 | *RAD51* in YEp13 | Wolf-Dietrich Heyer |
| pAM21 | *RAD51* in pUC13 | Lorraine Symington |
| AB076B | *RAD59* in pRS426 | Dennis Livingston |
| opb64b | *ELG1* N- terminus in pGBU9 | Lab stock |
| AB077B | *rad52-2* in pRS426 | Dennis Livingston |
| AB146 | *Rad52+Rad51* in pRS426 | Dennis Livingston |
| AB079 | *rad52-K43,44,253R* in pRS426 | This study |
| AB087 | *rad52-C180A* in pRS425 | This study |
| AB088 | *RAD52* in pRS425 | This study |
| MK3561 | *rad52-QDDD-AAAA* in pRS325 | (45) |
| MK3562 | *rad52-Δ327* in pRS325 | (45) |
| MK3563 | *rad52-R70A* in pRS325 | (45) |
| MK3563 | *rad52-R70A* in pRS325 | (45) |
| MK3564 | *rad52-L89F* in pRS325 | (45) |
| MK3354 | *RAD52* in pGAD424 | Giora Simchen |
| MK3353 | *RAD51* in pGAD424 | Giora Simchen |
| D2084 | *srs2(783-1174)* in pGBDC2 | Stefan Jentsch |
| D2089 | *srs2(783-1038)* in pGBD2 | Stefan Jentsch |
| AB169 | *srs2(783-1038)Δ875-902* in pGBD2 | This study |
| AB154 | *srs2(783-874)* in pGBT9 | This study |
| AB156 | *srs2(903-1038)* in pGBT9 | This study |
| AB135 | *SRS2* in pRS425 | This study |
| AB133 | *srs2-K41A* in pRS425 | This study |
| AB139 | *srs2Δ(875-902)* in pRS425 | This study |
| p195-*ELG1* | *ELG1-myc7HIS* in pYEplac195 | Lab stock |
| MK 1969 | *SMT3-pol30-k127r,k164r* in pGBT9 | (59) |

### DNA damage sensitivity serial dilution assay

Serial ten-fold dilutions of logarithmic yeast cells were spotted on fresh Synthetic Dextrose (SD)-complete (or SD lacking a specific amino acid to preserve the plasmid) plates with or without different concentrations of Methyl methane sulfonate (MMS)(Sigma) and incubated at 30°C for three days. MMS plates were freshly prepared, dried in a biological hood, and used the same day.

### Chromatin fractionation assay

50ml of a logarithmic culture was collected and washed with ddH_2_O PSB (20 mM Tris-HCl pH 7.4, 2 mM EDTA, 100 mM NaCl, 10 mM b-ME) and SB (1M Sorbitol, 20 mM Tris-HCl pH 7.4). Next cells were suspended in 1ml SB, 30 μl Zymolase 20T (20 mg/ml in SB) was added, and samples were incubated at 30 °C until spheroplasts were visible (around 1hrs). Spheroplasts were washed twice with SB and suspended in 500ml EBX (20 mM Tris-HCl pH 7.4, 100 mM NaCl, 0.25% Triton X-100, 15 mM β-ME + protease / phosphatase inhibitors). Triton X-100 was added to 0.5% final to lyse the outer cell membrane, and the samples kept on ice for 10 min with gentle mixing. Whole cell lysate (WCE) samples were taken and the rest of the lysate was layered over 1 ml NIB (20 mM Tris-HCl pH 7.4, 100 mM NaCl, 1.2 M Sucrose, 15mM β-ME + protease / phosphatase inhibitors). After centrifugation the cytoplasmic fraction was taken. The nuclear pellet was suspended in 500 μl EBX and Triton X-100 added to 1% final to lyse the nuclear membrane. The pellet was centrifuged and the chromatin was suspended in 50 μl Tris pH 8.0 for western analysis (Chromatin). For western blotting the following antibodies were used: PCNA antibody (1:1000,Abcam), histone H3 (1:5000, Abcam) RPS6(1:1000,Abcam).

### Yeast Two-hybrid assay

To detect two hybrid interactions, yeast strain PJ69 (58) was co-transformed with a *LEU2*-marked plasmid containing genes fused to the *GAL4* activating domain (pACT or pGAD424) and a plasmid containing genes fused to the *GAL4* DNA binding domain (pGBU9). Yeast cultures were grown in the respective selective media to retain the plasmids, with or without MMS. Cells were incubated for 3-5 days at 30°C.

## Acknowledgements

We thank Kara Bernstein, Wolf Heyer, Giora Simchen, Lorraine Symington, Hele Ulrich, Lumir Krejci and the late Stefan Jentsch for strains and plasmids, and past and presents members of the Kupiec lab for support, ideas and encouragement.

